# Bacterial purine metabolism modulates *C. elegans* development and stress tolerance via DAF-16 translocation

**DOI:** 10.1101/2023.09.17.558182

**Authors:** Min Feng, Baizhen Gao, L. Rene Garcia, Qing Sun

**Affiliations:** Department of Chemical Engineering, Texas A&M University, College Station, TX, USA; Department of Biology, Texas A&M University, College Station, TX, USA

**Author notes:** Correspondence: Qing Sun.

## Abstract

Purine homeostasis is crucial for cellular function and is a conserved metabolic network from prokaryotes to humans. While extensively studied in microorganisms like yeast and bacteria, the impact of perturbed dietary purine levels on animal development and balanced growth remains poorly understood. To investigate the mechanisms underlying this deficiency, we utilized *Caenorhabditis elegans* as the metazoan model. Through a high-throughput screening of an *E. coli* mutant library Keio collection, we identified 34 *E. coli* mutants that delay *C. elegans* development. Among these mutants, we found that *E. coli purE* gene is an essential genetic component that promotes host development in a dose-dependent manner. Additionally, we observed increased nuclear accumulation of the FoxO transcription factor DAF-16 when fed *E. coli purE-* mutants, suggesting the role of DAF-16 in response to nutrient, especially purine deficiency. RNA-seq analysis and phenotype assays revealed that worms fed the *E. coli purE* mutant exhibited elevated lifespan, thermotolerance, and pathogen resistance. These findings collectively suggest that perturbations in bacterial purine metabolism likely serve as a cue to promote development and activate the defense response in the nematode *C. elegans* through DAF-16 nuclear translocation.

## Introduction

Purine metabolism plays a critical role in maintaining cellular homeostasis and is conserved across various organisms, from prokaryotes to humans^1^. Purines serve as essential components of DNA and RNA, and their balanced regulation is crucial for cellular functions and overall organismal development^2^. While the mechanisms governing purine metabolism have been extensively studied in microorganisms, such as yeast and bacteria, limited knowledge exists regarding the effects of perturbed purine availability on animal development and the mechanism underlie balanced growth in animals.

*Caenorhabditis elegans*, a well-established model organism, offers unique advantages for investigating the intricate relationship between host development and dietary factors. As a bacterivore, *C. elegans* predominantly feeds on *Escherichia coli* strain OP50, which serves as the standard laboratory food source. Importantly, the genetic manipulability of bacteria allows for precise control over metabolite availability for *C. elegans*. In this study, we aim to understanding the role of microbial factors derived from *E. coli* in regulating host growth and development, with a specific focus on exploring the mechanisms underlying nutritional purine perturbation on *C. elegans* development.

The purine biosynthesis network is functionally conserved in *C. elegans*. Studies have revealed the importance of specific enzymes, such as adenylosuccinate lyase ADSL, in purine homeostasis. ADSL participates in both the purine *de novo* synthesis and recycling pathways. They are required for crucial developmental processes, including developmental timing, germline stem cell maintenance, and muscle integrity in *C. elegans*^*3*^.Whereas perturbations in purine metabolism are likely perceived as signals to promote defense mechanisms against epithelial infection in *C. elegans*^4^. High-purine diets, which caused uric acid accumulation, was proved to correlate with shortened lifespan in Drosophila^5^. Understanding the impact of dietary purine perturbation on animal development and health is of great importance, as it can provide valuable insights into the interplay between nutritional factors, metabolic pathways, and animal health.

In this study, we aim to expand our understanding of the complex interplay between nutritional factors, purine metabolism, and host development in *C. elegans*. By employing genetic and molecular techniques, we investigated the impact of purine availability, specifically focusing on the microbial factors derived from *E. coli*, on various aspects of *C. elegans* growth and development. The insights gained from this research will contribute to our broader understanding of the fundamental mechanisms underlying the regulation of growth and development in animals, with potential implications for human health and disease.

## Result

### Genome-wide screening reveals microbial factors impacting *C. elegans* development

To identify microbial factors that influence host development in *Caenorhabditis elegans*, we performed a systematic genome-wide screen using Keio collection, which contains 3985 single-gene knockout deletions of *E. coli* K12 strain BW25113^6^. Synchronized L1 worms were placed on individual *E. coli* mutants as the only food source, with the Keio parent strain BW25113 serving as the control. To capture variations in *C. elegans* growth when fed different bacterial mutants, we selected 48 hours as the time point to measure the worm’s body size; on average, worms fed wild-type *E. coli* BW25113 reach the young adult stage at this time point. In the primary screen, we examined the entire library and focused on deletion strains that exhibited an average variance in *C. elegans* body size greater than 10% compared to the control (Figure 1A). Interestingly, we did not observe any strains that increased the body size of *C. elegans*. Thus, deletion strains that resulted in a body size decrease greater than 10% were subjected to a secondary screening in triplicate. The body size of animals fed with these strains was measured using ImageJ software. Among the 3,983 *E. coli* mutants screened, 34 deletion strains were identified to cause a slower development of *C. elegans* compared to animals fed the wild-type *E. coli* (Figure 1B). These results indicate that the genetic composition of gut microbes has a significant impact on host development.

**Figure 1.**
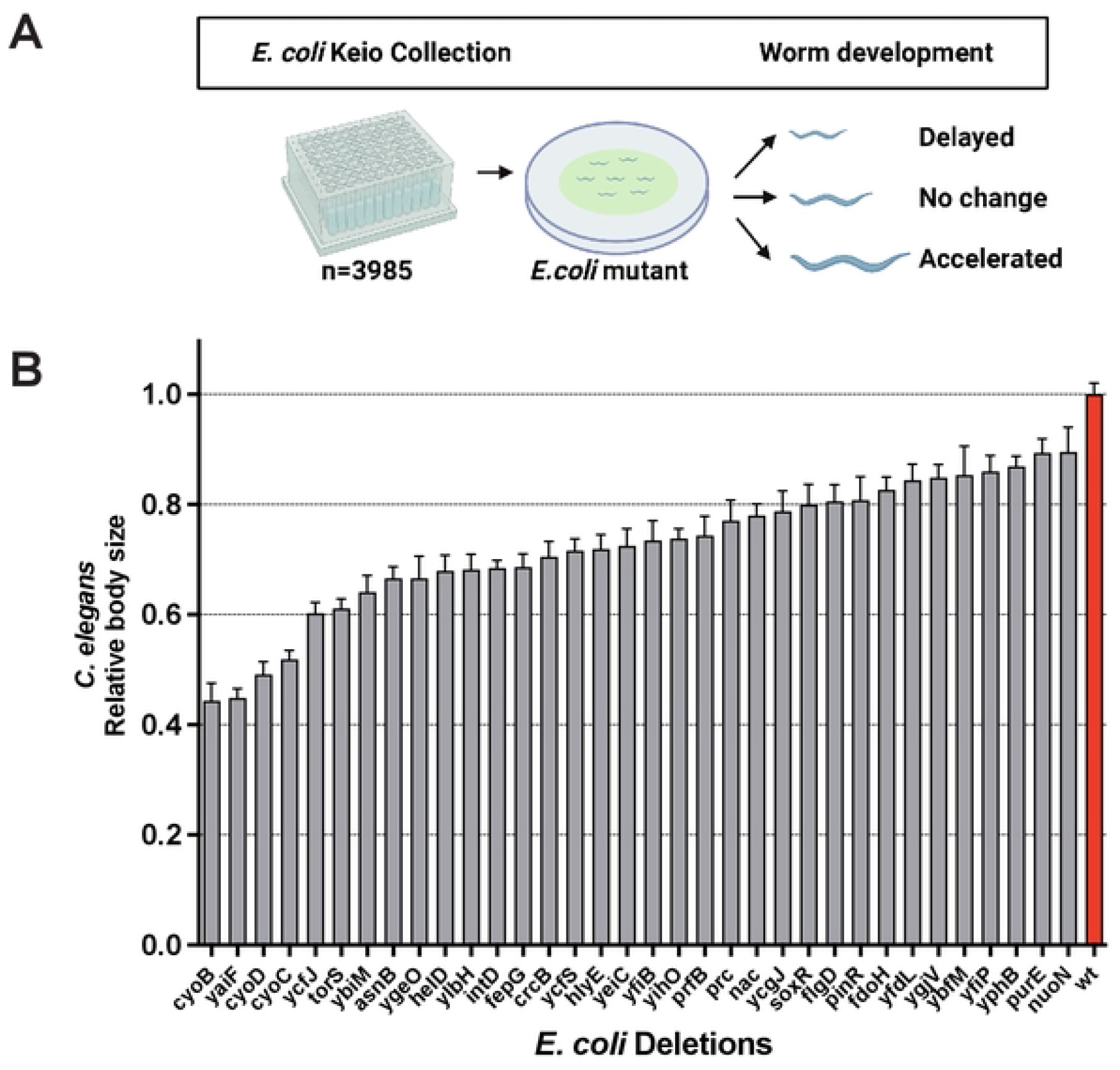
High-throughput screen of *E. coli* single gene deletions impacting *C. elegans* development. (A) Schematic representation of the genome-wide screens conducted. (B) Bar graph illustrating the effects of *E. coli* mutant strains from the Keio collection on worm development. The body size of worms was measured 2 days after synchronized L1 larvae were placed on the respective plates. Data are presented as mean ± SEM.

### The *E. coli purE* mutant reduces the developmental rate of *C. elegans*

In a previous publication, *E. coli* gene deletion mutants that hindered *C. elegans* development were identified, including genes involved in fatty acid metabolism and energy production^7^. Building upon these findings, our study aimed to discover additional *E. coli* genes affecting *C. elegans* development. Within all the identified genes, we focused on *E. coli purE* mutant that has deficient of an enzyme, *N*^*5*^-carboxyaminoimidazole ribonucleotide mutase, involved in purine nucleotide biosynthesis^8^. It catalyzes the reversible interconversion of *N*^*5*^-carboxyaminoimidazole ribonucleotide (*N*^*5*^-CAIR) and 5-amino-1-(5-phospho-D-ribosyl) imidazole-4-carboxylate (CAIR)^9^. Worms fed on the *purE* mutant exhibited slower development compared to those fed on wild-type *E. coli* (Figure 2A), suggesting insufficient nutrient from *E. coli purE* mutant for *C. elegans* development. To investigate if *E. coli purE* mutant serves as a poor nutritional source for *C. elegans* and limit growth/survival, we fed synchronized L1 worms a mixture of various ratios of wild-type *E. coli* and *purE* mutant. The body size of the worms was measured using ImageJ. The results demonstrated a decrease in worm size with increasing *purE* mutant ratio (Figure 2B), indicating that *E. coli purE* mutant was the reason for the reduction in worm development.

**Figure 2.**
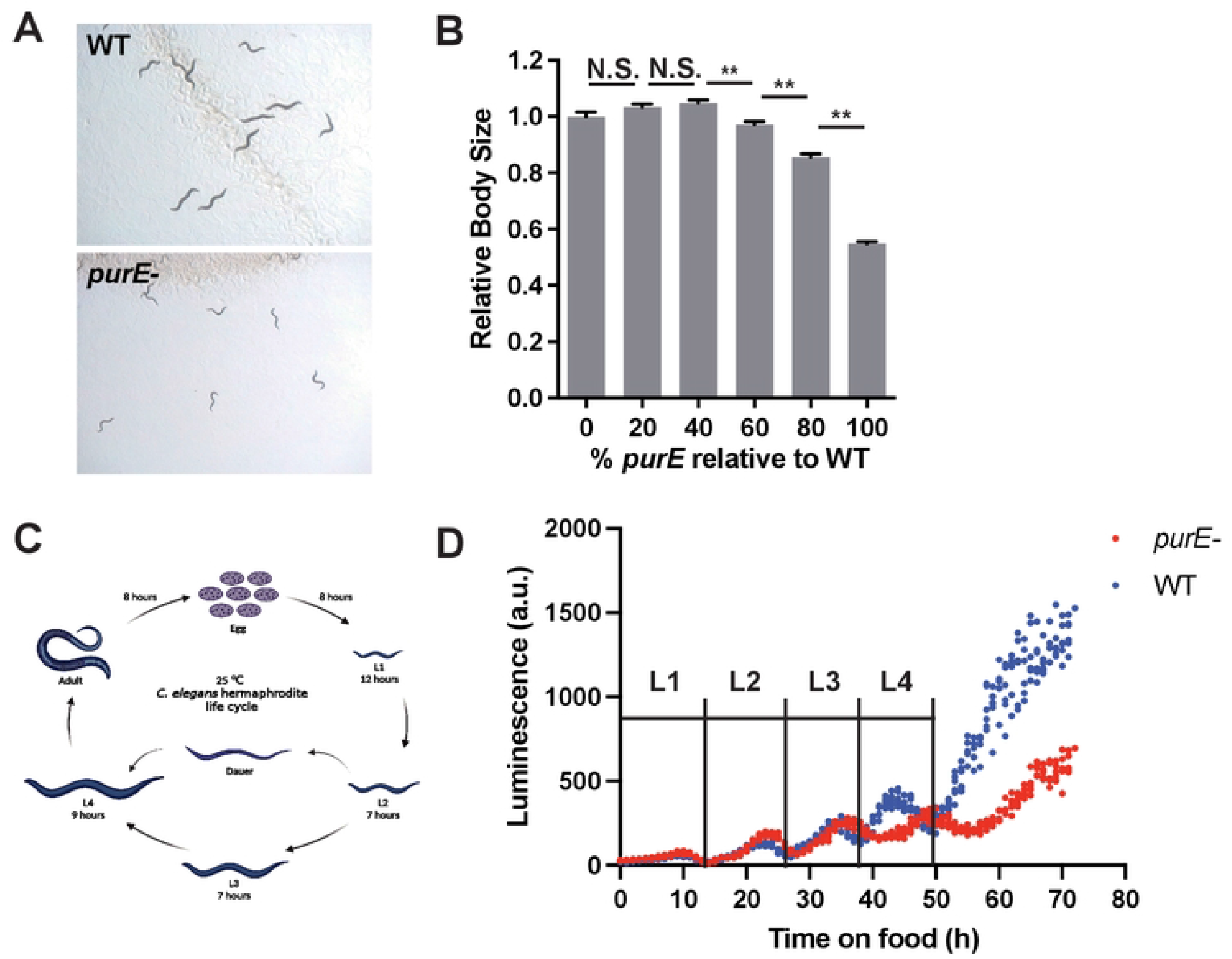
Impact of *E. coli* purine metabolism on host development. (A) Microscope images demonstrating the delayed development of worms fed with the *E. coli purE* mutant compared to those fed with wild-type *E. coli* BW25113. (B) Bar graph illustrating the growth of worms when fed with different ratios of PurE/WT bacterial mixture. Synchronized L1 worms were fed on the indicated ratio of purE/WT bacterial mixture, and the body size of worms was measured 2 days after synchronized L1 worms were placed on the plates. Data are presented as mean ± SEM. (C) Cartoon representation of the life cycle of C. elegans at 22°C. (D) Representative trace showing luminescentchanges during worm development.

The life cycle of *C. elegans* consists of four larval stages, L1, L2, L3, and L4 (Figure 2C). To assess whether the *E. coli purE* mutant caused a delay or arrest in *C. elegans* development, we incubated worms fed with the wild type or mutant strain for 72 hours. Remarkably, we observed that the worms eventually reached a size comparable to those grown on wild type *E. coli* and completed their development, with the majority of them initiating the production of offspring (Figure S1). Furthermore, luciferase-based development assays were performed to assess the growth rate of *C. elegans*, the expression of luciferase driven by promotors of *sur-5* gene that oscillate with the molts have been used as hallmarks of the developmental process^10^. The luminescence data revealed four distinct peaks, corresponding to the four larval stages in the *C. elegans* life cycle (Figure 2D). These results suggested that *C. elegans* fed with *purE* mutant had a slower rate compared with wild-type *E. coli* confirming that the observed slow growth was not due to developmental arrest.

### Role of *E. coli de novo* purine biosynthesis pathway in regulating *C. elegans* development

The primary purine nucleotide generation pathway in *E. coli* is the *de novo* purine nucleotide biosynthesis pathways^11^, where phosphoribosyl pyrophosphate (PRPP) serves as the initial substrate, leading to the formation of purine nucleotide IMP (Figure 3A). To explore the regulatory role of this bacterial pathway in worm development, we fed *C. elegans* with various *E. coli* mutants as food and assessed worm size as indication of worm growth. We found that worm development was significantly delayed when fed with *E. coli* mutants *purC, purE*, and *purF*, which were the key genes in *de novo* purine biosynthesis pathway (Figure 3B). These results indicate the crucial involvement of the *E. coli de novo* purine biosynthesis pathway in promoting *C. elegans* growth.

**Figure 3.**
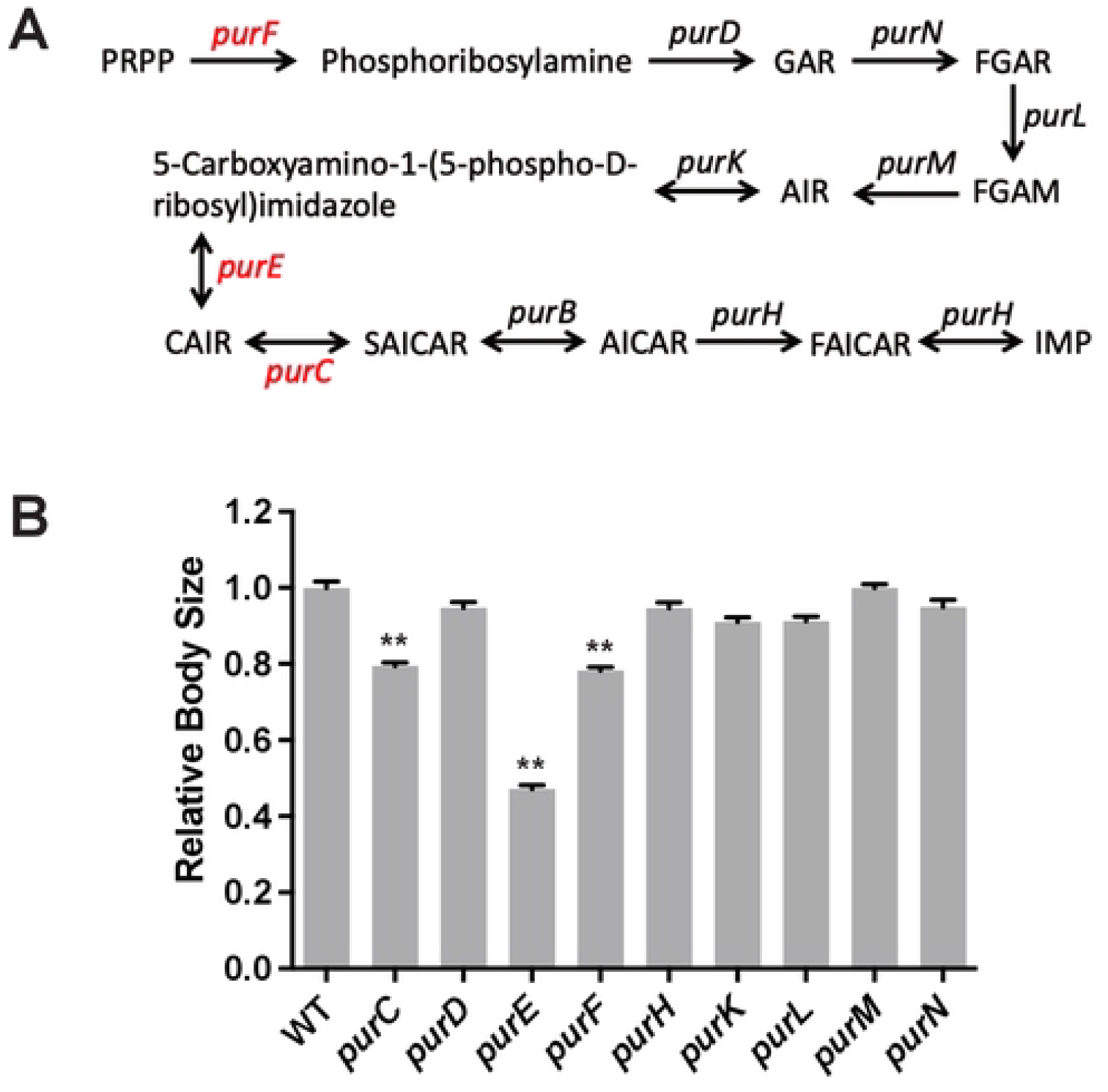
Impact of perturbations in *E. coli* de novo purine nucleotide biosynthesis pathway on *C. elegans* development. (A) Homologs of the purine nucleotide biosynthesis pathway in E. coli. Abbreviations: PRPP, 5-phosphoribosyl diphosphate; GAR, glycinamide ribonucleotide; FGAR, N -formylglycinamide ribonucleotide; FGAM, 5-phosphoribosylformylglycinamidine; AIR, aminoimidazole ribotide; CAIR, 5-phosphoribosyl-4-carboxy-5-aminoimidazole; SAICAR, 5-phosphoribosyl-4-(N -succinocarboxamide)-5-aminoimidazole; AICAR, 5-aminoimidazole-4-carboxamide ribotide; FIACAR, 5-phosphoribosyl-5-formamido-4-imidazolecarboxamide. (B) Bar graph illustrating the growth of worms when fed with E. coli mutants involved in the purine nucleotide biosynthesis pathway. Synchronized L1 worms were fed on the indicated E. coli mutants, and the body size of worms was measured 2 days after synchronized L1 worms were placed on the plates. Data are presented as mean ± SEM.

### Bacterial *purE* mutant induced host DAF-16 nuclear translocation

To investigate the molecular mechanisms underlying the growth effects of *E. coli purE* mutant, we examined its interactions with several well-known cellular processes involved in host growth and development. Specifically, we focused on SEK-1, a protein involved in the mitogen-activated protein kinase (MAPK) signaling pathway regulating cell division and differentiation; EAT-2, which encodes a nicotinic acetylcholine receptor subunit involved in feeding behavior and nutrient uptake; DAF-16, a transcription factor downstream of the insulin/IGF-1 signaling pathway regulating metabolism and stress response; AKT-1, a serine/threonine kinase controlling cell growth and survival; and SKN-1, a transcription factor involved in oxidative stress response and detoxification. To determine the cellular process’ effects on host developmental delay caused by *E. coli purE* mutant, we investigated the development of *C. elegans* mutants *sek-1(km4), eat-2(ad465), daf-16(mgDf50), akt-1(ok525)*, and *skn-1(zu135)* grown on the *E. coli purE* mutant for 48 hours at RT. Our screening revealed that among all the *C. elegans* mutants, *E. coli purE* mutant did not significantly alter the body size of *C. elegans daf-16* mutant, nor enhance any defects (Figure 4A), suggesting *C. elegans* insulin/IGF-1 signaling pathway as a regulatory mechanism influencing development.

**Figure 4.**
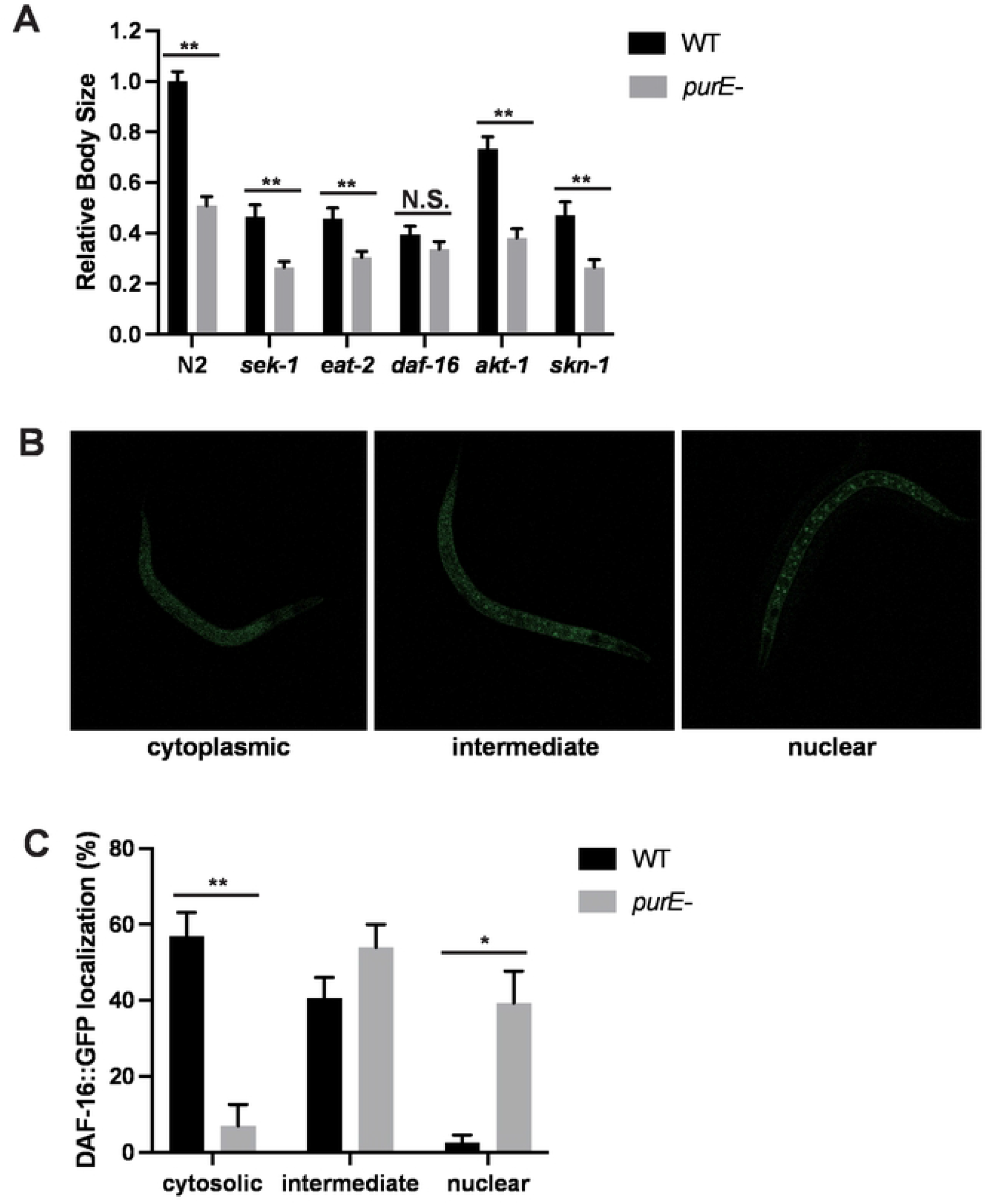
Essential role of DAF-16 in bacterial *purE-* mediated host development. (A) Bar graph illustrating the growth of wild-type N2, *sek-1, eat-2, daf-16, akt-1*, and *skn-1* mutant worms when fed on *E. coli* parent strain BW25113 or *purE* mutant strain. Synchronized L1 worms of different genotypes were fed on either wild-type *E. coli* strain BW25113 or *purE* mutant strain, and the body size of the worms was measured 48 hours after L1 stage placement on the plates. (B) Microscope images displaying representative images of the transgenic strain TJ356 with cytosolic, intermediate, and nuclear localization of DAF-16::GFP. (C) Bar graph demonstrating the activation of DAF-16::GFP nuclear translocation in worms fed on the *E. coli purE* mutant. Data are presented as mean ± SEM, with n > 200 in three independent experiments; ** denotes P < 0.01, * denotes P < 0.05.

DAF-16, a FoxO transcription factor that is regulated by insulin-signaling pathway, responds rapidly to environmental stimuli by translocation from cytoplasm to nucleus, and subsequently activates transcription^12,13^. To investigate the effect of *E. coli purE* mutant on this signaling pathway, we examined whether the mutant bacteria induced nuclear translocation of DAF-16 using a transgenic strain expressing fusion protein DAF-16::GFP (TJ356). In Figure 4B, three different statuses of DAF-16 localization were shown: cytosolic (left), intermediate (center), and nuclear (right). When *C. elegans* were fed with the *E. coli purE* mutant, we observed a shift in DAF-16 localization from cytosol to nucleus (Figure 4C), suggesting that delayed worm development caused by *E. coli purE* mutant could be attributed to the nuclear localization of DAF-16.

### RNA-seq reveals the effect of *E. coli purE* mutant on host iron homeostasis and fat storage

To investigate the signaling mechanism underlying the developmental delay of *C. elegans* fed on *E. coli purE* mutant, we performed RNA sequencing to identify host pathways regulated by *E. coli purE* activity. Specifically, we compared gene expression profiles between worms fed the *E. coli purE* mutant diet and wild-type *E. coli* BW25113 diet. To mitigate the impact of developmental stage differences, worms were fed either the wild-type *E. coli* BW25113 or *E. coli purE* mutant for 24 hours and then switched to a wild-type *E. coli* BW25113 or *E. coli purE* mutant diet for 2 hours prior to RNA extraction and RNA-seq analysis (Figure 5A). Employing a fold-change difference of ≥1 and a P-value < 0.05, we identified 45 genes with increased expression and 79 genes with reduced expression in worms fed the *E. coli purE* mutant compared to wild-type *E. coli* (Group 2 vs. Group 1), as well as 91 genes with increased expression and 289 genes with reduced expression upon diet switch (Group 4 vs. Group 3) (Figure 5B) (Table S1). Furthermore, qPCR analysis confirmed the relative mRNA level changes of several representative upregulated genes (*nhr-202, argk-1, cth-1*, and *cyp-35d1*) and downregulated genes (*y41c4a-32* and *cpt-4*) compared to the internal reference gene *act-1* (Figure 5C).

**Figure 5.**
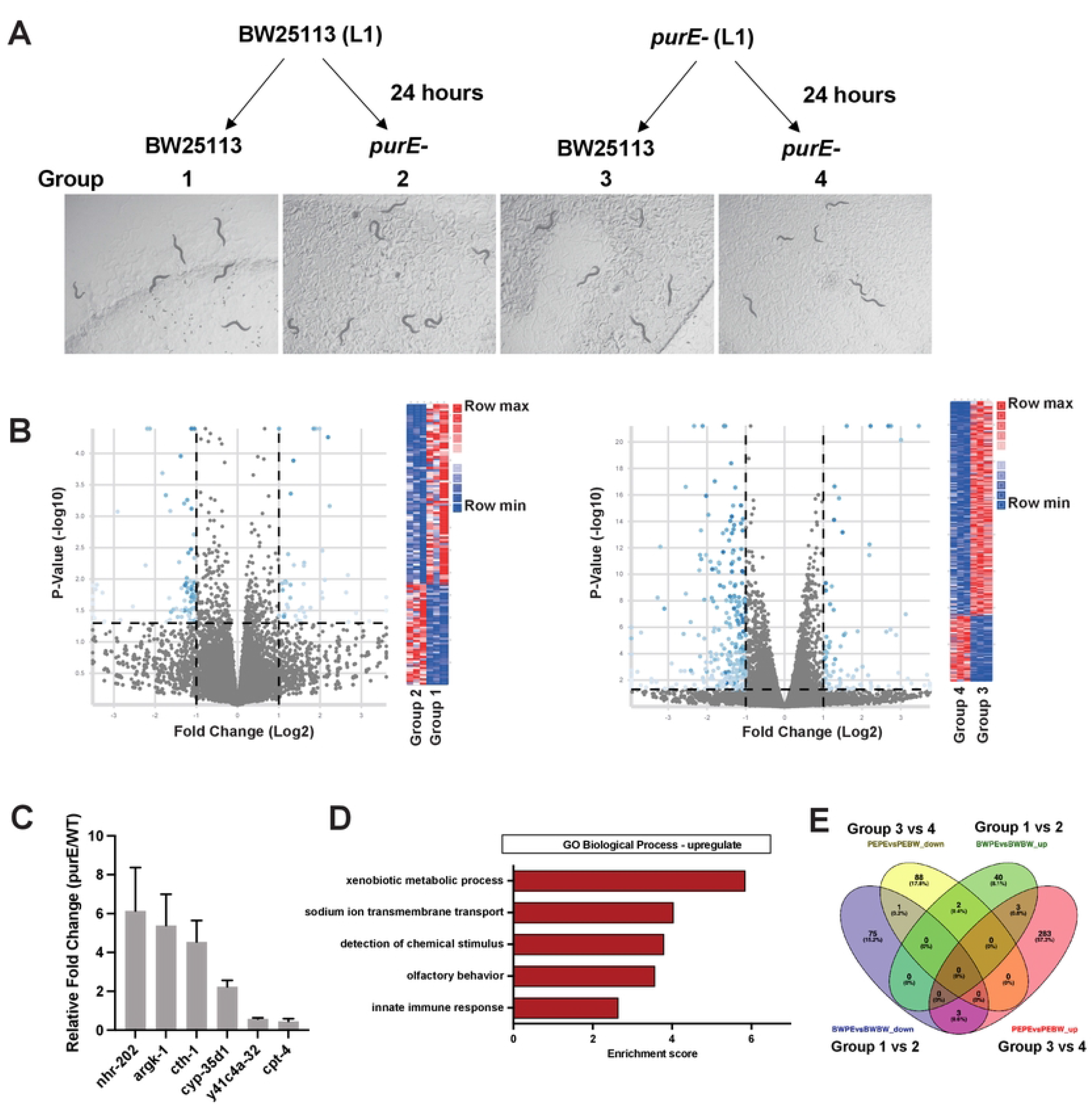
RNA-seq analysis reveals enhanced immune resistance in worms fed with *E. coli purE* mutant. (A) Workflow illustrating the preparation of worm samples and representative images displaying worms developed under different experimental conditions. (B) Volcano plots and heat map displaying the fold-change and P-value of differentially expressed genes (DEGs) observed in the RNA-seq comparison of Group 2 versus Group 1. (C) RT-qPCR results confirming the upregulated or downregulated genes consistent with the RNA-seq findings. (D) GO analysis revealing enrichment in several biological processes. (E) Venn diagram demonstrating the common upregulated genes upon diet switch.

Functional enrichment analysis using Gene Ontology (GO) revealed that the upregulated genes were primarily associated with xenobiotic metabolic processes, sodium ion transmembrane transport, chemical stimulus detection, olfactory behavior, and immune response (Figure 5D), consistent with the crucial role of iron homeostasis in animal development^7^. Notably, the Venn diagram indicated common upregulated genes (*fbxa-48, cyp-35D1*, and *hrg-3*) and a common downregulated gene (*fat-7*), suggesting the key role of fat-7 in regulating fat storage in response to this dietary change. Collectively, these results highlight the fundamental role of iron homeostasis in regulating host development under conditions of purine perturbation, with *fat-7* emerging as a potential regulator of decreased fat storage.

### Bacterial *purE* induces host innate immune response and stress resistance

Since the effects of purine metabolism perturbation in the diet had not been previously characterized in *C. elegans*, we further explored additional phenotypes including host immune response and stress resistance. Firstly, we assessed the thermotolerance of worms fed with *purE* mutant. To minimize potential age-related differences, synchronized L1 worms were cultured at a room temperature of 21.5°C for 48 hours on a wild-type *E. coli* diet or 55 hours on a *purE* mutant diet, ensuring the worms at the same developmental stage. Then adult worms raised on different diets were subjected to a 2-hour incubation at 37°C, followed by recovery at room temperature for 24 hours, and their survival was measured. Results revealed increased thermotolerance of worms fed with *purE* mutant compared to wild-type *E. coli* (Figure 6A). Secondly, we investigated the pathogen resistance phenotypes of worms fed with *purE* mutant using slow-killing assay. We found reduced susceptibility to the Gram-negative bacterial pathogen *Pseudomonas aeruginosa* strain PA14 in worms fed the *purE* mutant (Figure 6B). Additionally, we explored the lifespan of worms fed the *purE* mutant and found a significant increase compared to those fed wild-type *E. coli* (Figure 6C), consistent with findings from a systematic screening of microbial genetic variations associated with longevity promotion^14^. Taken together, these results clearly demonstrate that bacterial purine metabolism impacts host thermotolerance, pathogen resistance, and lifespan.

**Figure 6.**
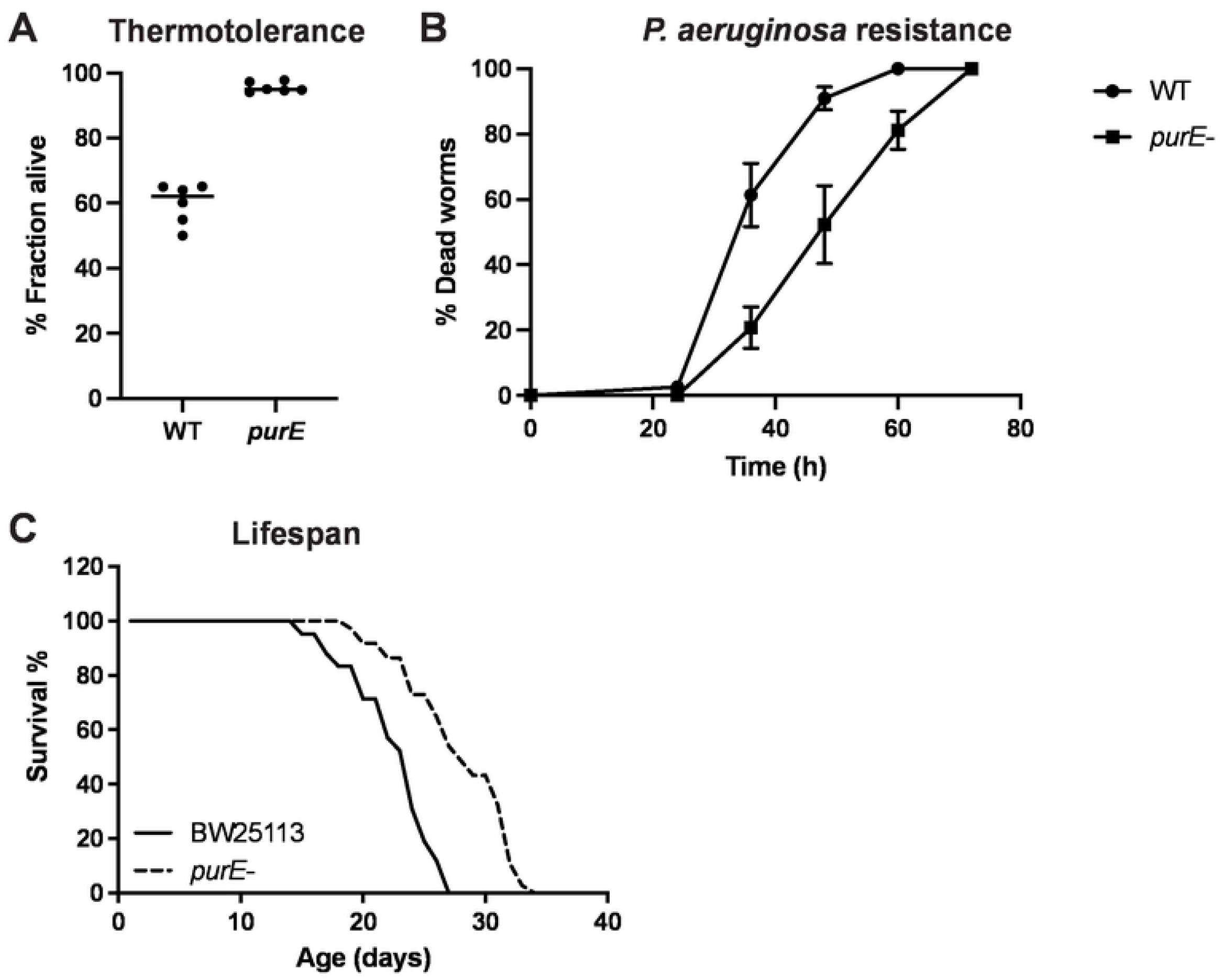
Lifespan, thermotolerance, and *Pseudomonas aeruginosa* resistance phenotypes of worms fed the *E. coli purE* mutant. (A) Lifespan analysis of worms fed on wild-type *E. coli* and the *purE* mutant. (B) Survival of worms fed on wild-type *E. coli* and the *purE* mutant following a 2-hour heat shock at 37°C, measured 24 hours later. (C) Slow killing assay with *P. aeruginosa* strain PA14 to assess resistance in worms fed on wild-type *E. coli* and the *purE* mutant.

## Discussion

In this study, we investigated the influence of nutritional purine perturbation on *C. elegans* growth and development. Especially, we used *E. coli* as model microbial system to study the role of nutritional deficiency in modulating host responses. Our results confirm the disruption of *E. coli purE* gene, involved in purine biosynthesis, leads to a delay in *C. elegans* development, indicating the importance of purine metabolism in host physiology. Among all the *E. coli* deletions involved in *de novo* purine biosynthesis pathway, the *purE* mutant, which affects the N5-CAIR conversion, might lead to a more critical disruption in the downstream biosynthesis of purine nucleotides, impacting crucial cellular processes required for worm growth. On the other hand, the *purC* and *purF* mutants might have slightly less pronounced effects (Figure 3B), as they are involved in different stages of the de novo purine biosynthesis pathway. As for the lack of obvious effects seen with *purD, purH, purK, purL, purM*, and *purN* mutants (Figure 3B), it could be due to compensatory mechanisms in the purine biosynthesis network by alternative pathways or redundant enzymatic activities. In the context of the observed worm growth, these mutants might not be critical nodes that would lead to significant growth alterations when disrupted.

Moreover, *C. elegans* fed the *purE* mutant displayed notable enhancements in thermotolerance, heightened resistance to pathogens, and increased lifespan in comparison to those fed wild-type *E. coli* (Figure 6). Remarkably, these phenotypic outcomes similar to the effects of caloric restriction. Intriguingly, our investigations into EAT-2, a gene encoding a nicotinic acetylcholine receptor subunit linked to feeding behavior and nutrient uptake, revealed intriguing distinctions. When *eat-2* mutant *C. elegans* were fed with wild-type *E. coli* or the *purE* mutant for 48 hours, a substantial size differential persisted (Figure 4A). This observation suggests that the mechanisms underlying the phenotypic shifts induced by bacterial *purE* alterations differ from those triggered by caloric restriction.

Monounsaturated fatty acids play a vital role in forming membrane and storage lipids, and their synthesis hinges on the conversion of saturated fatty acids into unsaturated fatty acids through enzymes like stearoyl-CoA desaturases (SCDs)^15^. Interestingly, prior research has showcased how SCDs respond to shifting environmental circumstances in mouse models. For instance, mice exhibit reduced SCD expression when presented with a diet rich in unsaturated fatty acids, this regulation extends to dietary carbohydrates and hormones like insulin and leptin^16^. In *C. elegans*, one of the SCD-encoding gene is *fat-7*. Our study revealed *fat-7* emerged as a compelling candidate for orchestrating reduced fat storage under conditions of purine perturbation (Figure 5E). This finding adds a layer of complexity to the regulation of lipid metabolism, suggesting that *fat-7* might serve as a responsive mediator in the intricate interplay between this specific nutritional cues and lipid homeostasis.

Iron homeostasis is crucial for biological processes of all life forms^17^. Nutritional imbalances affecting iron levels pose a significant threat to global health^18^. In our study, the RNA-seq analysis unveiled extensive alterations in host gene expression, notably within enriched functional categories related to sodium ion transmembrane transport (Figure 5D). The intricate interplay between the genetic composition of gut bacteria, which produces a range of purine synthesis enzymes, has the potential to exert a profound influence on maintaining iron homeostasis. The striking impact of bacterial purine homeostasis on both iron transmembrane transport and animal growth indicates that disruptions in microbial composition could substantially contribute to human iron imbalances. This novel insight carries significant implications, suggesting the potential application of such disruptions as a means to address and maintain iron homeostasis, thereby promoting overall health.

These findings emphasize the crucial role of purine metabolism in *C. elegans* development and highlight the significant impact of gut microbial factors on host physiology through modulation of purine availability. The study provides valuable insights into the complex interplay between microbial purine metabolism, host signaling pathways, and phenotypic traits. Ultimately, these findings could have implications for understanding host-microbiota interactions and their potential relevance to human health.

## Methods

### *C. elegans* and bacterial strains

Worms were maintained by standard protocols following worm book. Bristol N2 was used as the wild type strain. PE254 ((*feIs4* [P*sur-5*:luc+::gfp; *rol-6* (su1006)]V) was used for the developmental assay. TJ356 [zIs356 IV (P*daf-16*:*daf-16*::gfp; *rol-6*)]. All experiments were performed with synchronized L1 hermaphrodites that incubated for 48 hours on nematode growth media (NGM) at room temperature. *E. coli* strain OP50 and wild type *E. coli* BW25113 were grown at 37°C in Lysogeny Broth (LB). *E. coli* deletion mutants (Baba et al., 2006) were grown at 37°C in LB with 50 μg/mL kanamycin. Overnight cultures were seeded on NGM plates and dried for 1 hour at room temperature before experiment.

### *E. coli* deletion library screen for changes in *C. elegans* developmental rate

Bacteria were grown overnight at 37°C in LB medium with 50 μg/mL kanamycin in 96-well deep plates. 150 μL of the overnight culture was seeded onto 35 mm NGM plate and dried at room temperature prior to the addition of *C. elegans*. Wild type N2 worms were fed the parental *E. coli* BW25113 strain for at least two generations prior to screening. Embryos were harvested from gravid worms using bleach solution and incubated overnight in M9 buffer with gentle shaking at room temperature to get synchronized L1 worms. Approximately 100 synchronized L1 worms were placed in each 35 mm NGM plates containing individual *E. coli* single-gene deletion strains and allowed to grow for 48 hours at room temperature. Bright-field images of worms in each well were collected using Leica S9i imaging system, and body size was determined using ImageJ. Each *E. coli* single gene deletion strain was screened once and mutants that cause obvious worm body size change were retested in triplicate in 35 mm plates. Bacterial mutants that caused at least a 10% reduction in animal body size in each of the three retest experiments were considered as positive hits.

### Luciferase developmental rate assay

Luciferase developmental rate assays were performed as described (Olmedo et al., 2015). Briefly, 5 arrested L1 animals were transferred into one well of a 96-well plate containing 100 μl of S-basal medium with 200 mM D-Luciferin and simultaneously resumed development by adding 100 μl of S-basal with bacteria. Plates were sealed with a gas-permeable cover (Breathe Easier, Diversified Biotech) and luminescence was measured using BioTek Synergy H1 for 1 sec, at 5-min intervals.

### Intracellular localization of DAF-16::GFP

Transgenic strain TJ356 [zIs356 IV (P*daf-16*-daf:16::gfp; rol-6)] was used to detect the intracellular localization of GFP tagged DAF-16 protein. Synchronized L1 worms were placed onto the NGM plates containing wild-type E. coli BW25113 or *E. coli purE* mutant and incubated for 48 h at room temperature. Subsequently, worms were placed on the glass slides, covered with cover slips and the cellular localization of DAF-16::GFP was detected by confocal microscopy. The experiment was repeated three times.

### RNA-seq

Synchronized L1 stage *C. elegans* worms were grown on NGM agar plates seeded with *E. coli* OP50 bacteria. After reaching the desired developmental stage, worms were transferred to NGM agar plates with either wild-type *E. coli* BW25113 or the *E. coli purE* mutant. Following a 24-hour feeding period, worms underwent a 2-hour diet switch. Worms were collected, and total RNA was extracted. The RNA was treated with DNase to remove genomic DNA contamination. The quality and quantity of RNA were assessed and used for library preparation. Libraries were then sequenced on an Illumina HiSeq 2000 sequencing machine using paired-end sequencing with a read length of 100 nucleotides. Adaptor sequences and low-quality reads were removed using Trimmomatic, The RNA-Seq reads were then mapped to the C. elegans genome using STAR 2.5.3a with default settings. Transcript abundance, measured as read counts per gene, was extracted using HTSeq. Differential gene expression analysis was performed using Featurecounts. DAVID bioinformatic database was used for gene enrichment analysis.

### RT-qPCR

RNA was extracted using TRIzol reagent (Invitrogen), according to the manufacturer’s instructions, and converted to cDNA using the iScript (Bio-Rad) cDNA synthesis kit. qPCR was performed using iQ SYBR green supermix (Bio-Rad) and various gene specific primers (Table S2) on a BioRad CFX Connect real-time system. The relative expression levels of target genes were calculated using a reference gene. Three experimental replicates were performed with one biological replicate for each condition. Unpaired, one-tailed student t-test was used. A p-value < 0.05 was considered significantly different.

### Lifespan assay

For lifespan assays, ∼50 synchronized L1 worms/plate per strain were plated onto 3.5-NGM plates seeded with BW25113 or *purE* mutant. Plates were incubated at room temperature.

After 48 hours, 30 adults per strain were scored in triplicate and live animals were transferred to fresh plates. Live worms were transferred every 24 hours until death or until progeny production stopped. Survival was measured every 24 hours and worms that did not respond to touch were scored as dead. Three experimental replicates were performed.

### Thermotolerance assay

Synchronized L1 worms were cultured at a room temperature of 21.5°C for 48 hours on a wild-type *E. coli* diet or 55 hours on a *purE* mutant diet, ensuring the worms at the same developmental stage. Then adult worms raised on different diets were subjected to a 2-hour incubation at 37°C, followed by recovery at room temperature for 24 hours, and their survival was measured. During both the heat shock and recovery, plates were placed in a single layer (plates were not stacked on top of each other). Worms were defined as dead by lack of movement on the plate, lack of pharyngeal pumping and lack of response to touch. For each replicate, three plates containing 30 worms were scored per genotype. Three experimental replicates were performed.

### Slow-killing assay

*P. aeruginosa* (PA14) killing assays were performed as previously described^19^. Three experimental replicates were performed.

### Statistical analysis

The statistical analyses were conducted using GraphPad Prism and Excel. The results are presented as the mean ± SEM, and the data were evaluated using an unpaired two-tailed Student’s t-test.

## Acknowledgments

This work was supported by the Texas A&M Startup grant from TEES and Department of Chemical Engineering to Q.S.

---

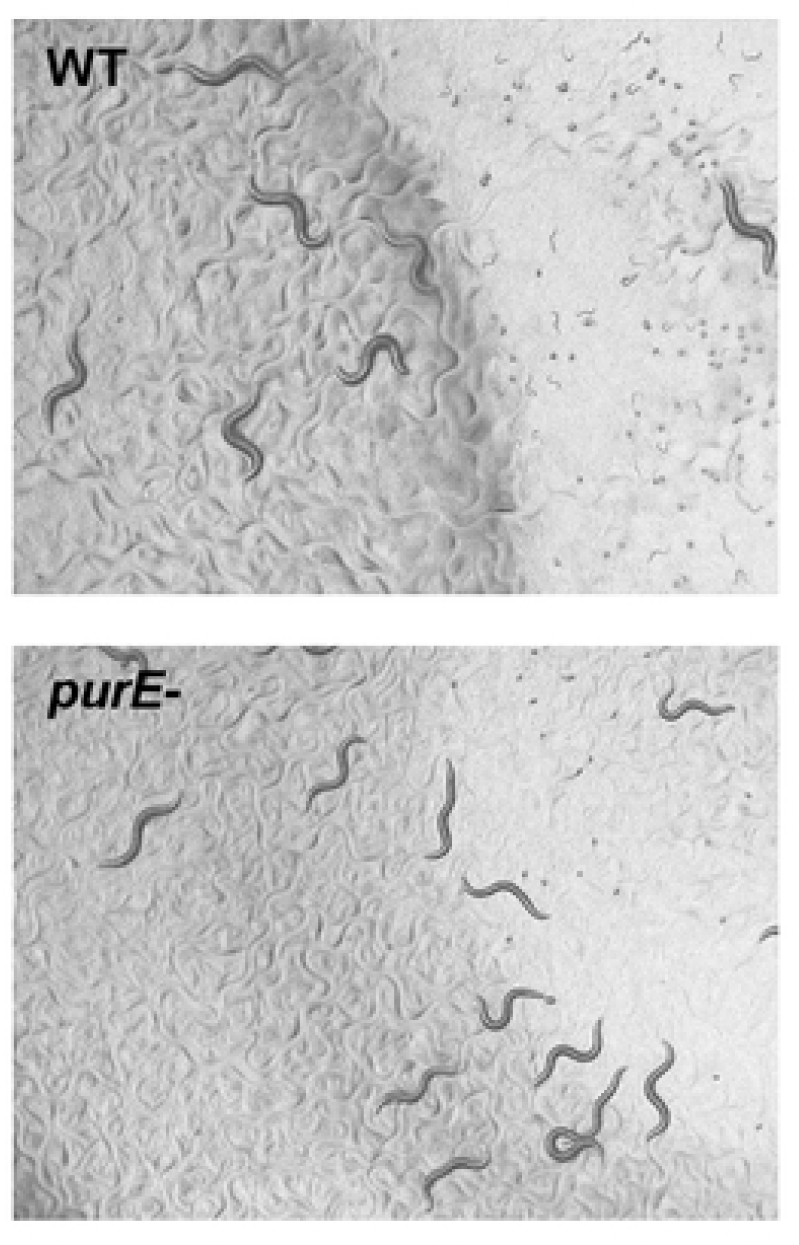

## Reference

1. Rolfes, R.J. (2006). Regulation of purine nucleotide biosynthesis: in yeast and beyond. Biochemical Society Transactions 34, 786–790. 10.1042/BST0340786.

2. Pedley, A.M., and Benkovic, S.J. (2017). A New View into the Regulation of Purine Metabolism – The Purinosome. Trends Biochem Sci 42, 141–154. 10.1016/j.tibs.2016.09.009.

3. Marsac, R., Pinson, B., Saint-Marc, C., Olmedo, M., Artal-Sanz, M., Daignan-Fornier, B., and Gomes, J.-E. (2019). Purine Homeostasis Is Necessary for Developmental Timing, Germline Maintenance and Muscle Integrity in Caenorhabditis elegans. Genetics 211, 1297–1313. 10.1534/genetics.118.301062.

4. Tecle, E., Chhan, C.B., Franklin, L., Underwood, R.S., Hanna-Rose, W., and Troemel, E.R. (2021). The purine nucleoside phosphorylase pnp-1 regulates epithelial cell resistance to infection in C. elegans. PLoS pathogens 17, e1009350.

5. van Dam, E., van Leeuwen, L.A.G., dos Santos, E., James, J., Best, L., Lennicke, C., Vincent, A.J., Marinos, G., Foley, A., Buricova, M., et al. (2020). Sugar-Induced Obesity and Insulin Resistance Are Uncoupled from Shortened Survival in Drosophila. Cell Metabolism 31, 710–725.e7. 10.1016/j.cmet.2020.02.016.

6. Baba, T., Ara, T., Hasegawa, M., Takai, Y., Okumura, Y., Baba, M., Datsenko, K.A., Tomita, M., Wanner, B.L., and Mori, H. (2006). Construction of Escherichia coli K-12 in-frame, singlegene knockout mutants: the Keio collection. Mol Syst Biol 2, 2006.0008. 10.1038/msb4100050.

7. Zhang, J., Li, X., Olmedo, M., Holdorf, A.D., Shang, Y., Artal-Sanz, M., Yilmaz, L.S., and Walhout, A.J.M. (2019). A Delicate Balance between Bacterial Iron and Reactive Oxygen Species Supports Optimal C. elegans Development. Cell Host &Microbe 26, 400–411.e3. 10.1016/j.chom.2019.07.010.

8. Mathews, I.I., Kappock, T.J., Stubbe, J., and Ealick, S.E. (1999). Crystal structure of Escherichia coli PurE, an unusual mutase in the purine biosynthetic pathway. Structure 7, 1395–1406. 10.1016/S0969-2126(00)80029-5.

9. Hoskins, A.A., Morar, M., Kappock, T.J., Mathews, I.I., Zaugg, J.B., Barder, T.E., Peng, P., Okamoto, A., Ealick, S.E., and Stubbe, J. (2007). N5-CAIR mutase: role of a CO2 binding site and substrate movement in catalysis. Biochemistry 46, 2842–2855. 10.1021/bi602436g.

10. Lagido, C., Pettitt, J., Flett, A., and Glover, L.A. (2008). Bridging the phenotypic gap: Real-time assessment of mitochondrial function and metabolism of the nematode Caenorhabditis elegans. BMC Physiology 8, 7. 10.1186/1472-6793-8-7.

11. Gedeon, A., Karimova, G., Ayoub, N., Dairou, J., Giai Gianetto, Q., Vichier-Guerre, S., Vidalain, P.-O., Ladant, D., and Munier-Lehmann, H. (2023). Interaction network among de novo purine nucleotide biosynthesis enzymes in Escherichia coli. The FEBS Journal 290, 3165–3184. 10.1111/febs.16746.

12. Oh, S.W., Mukhopadhyay, A., Svrzikapa, N., Jiang, F., Davis, R.J., and Tissenbaum, H.A. (2005). JNK regulates lifespan in Caenorhabditis elegans by modulating nuclear translocation of forkhead transcription factor/DAF-16. Proc Natl Acad Sci U S A 102, 4494–4499. 10.1073/pnas.0500749102.

13. Henderson, S.T., and Johnson, T.E. (2001). daf-16 integrates developmental and environmental inputs to mediate aging in the nematode Caenorhabditis elegans. Curr Biol 11, 1975–1980. 10.1016/s0960-9822(01)00594-2.

14. Han, B., Sivaramakrishnan, P., Lin, C.-C.J., Neve, I.A.A., He, J., Tay, L.W.R., Sowa, J.N., Sizovs, A., Du, G., Wang, J., et al. (2017). Microbial Genetic Composition Tunes Host Longevity. Cell 169, 1249–1262.e13. 10.1016/j.cell.2017.05.036.

15. Brock, T.J., Browse, J., and Watts, J.L. (2007). Fatty Acid Desaturation and the Regulation of Adiposity in Caenorhabditis elegans. Genetics 176, 865–875. 10.1534/genetics.107.071860.

16. Ntambi, J.M., and Miyazaki, M. (2004). Regulation of stearoyl-CoA desaturases and role in metabolism. Prog Lipid Res 43, 91–104. 10.1016/s0163-7827(03)00039-0.

17. Nakamura, T., Naguro, I., and Ichijo, H. (2019). Iron homeostasis and iron-regulated ROS in cell death, senescence and human diseases. Biochimica et Biophysica Acta (BBA) - General Subjects 1863, 1398–1409. 10.1016/j.bbagen.2019.06.010.

18. Kassebaum, N.J., Jasrasaria, R., Naghavi, M., Wulf, S.K., Johns, N., Lozano, R., Regan, M., Weatherall, D., Chou, D.P., Eisele, T.P., et al. (2014). A systematic analysis of global anemia burden from 1990 to 2010. Blood 123, 615–624. 10.1182/blood-2013-06-508325.

19. Powell, J.R., and Ausubel, F.M. (2008). Models of Caenorhabditis elegans Infection by Bacterial and Fungal Pathogens. In Innate Immunity Methods in Molecular Biology™., J. Ewbank and E. Vivier, eds. (Humana Press), pp. 403–427. 10.1007/978-1-59745-570-1_24.

